# Redox-Activated Proton Transfer through a Redundant Network in the Q_*o*_ Site of Cytochrome *bc*_1_

**DOI:** 10.1101/2024.11.22.624873

**Authors:** Guilherme M. Arantes

## Abstract

Proton translocation catalyzed by cytochrome *bc*_1_ (respiratory complex III) during coenzyme-Q redox cycling is a critical bioenergetic process, yet its detailed molecular mechanism remains incompletely understood. Even the specific groups mediating proton transfer following coenzyme-Q oxidation have not been established. In this study, the energetics of proton transfer through multiple proton-conducting wires recently identified in the Q_*o*_ site was investigated across various reactant redox states using hybrid QM/MM simulations and a specialized reaction coordinate. Key reactive groups and proton transfer mechanisms were characterized, confirming the propionate-A group of heme *b*_*L*_ as a plausible proton acceptor. Upon quinol oxidation, a Grotthuss hopping mechanism is activated, facilitating proton transfer along three distinct pathways with comparable energetics. These pathways function redundantly, forming a robust proton-conducting network. A highly conserved tyrosine residue (Y147 in *R. sphaeroides* numbering) was found to be essential for complete proton transfer, whereas participation of H276 and D278 does not appear energetically feasible. Bioenergetic analyses exclude charged closed-shell species as likely intermediates and propose a reaction sequence for quinol oxidation proceeding as QH_2_ → QH^*•*^ → Q^0^, either via coupled proton-electron transfers or stepwise mechanisms involving open-shell intermediates. These findings elucidate mechanistic details of the Q-cycle and improve our understanding of the catalytic reactions supporting coenzyme-Q redox cycling in respiratory complexes.

## Introduction

Cytochrome bc_1_, also known as respiratory complex III, is fundamental for cellular respiration and photosynthesis.^1^ It catalyzes the Q-cycle,^2–4^ where the membrane-soluble, two-electron carrier coenzyme-Q (here generically named Q) undergoes redox reactions that sustain the electron transport chain and translocate protons across the membrane. In addition to its primary role in energy transduction, the bc_1_ complex can generate reactive oxygen species, linking it to metabolic dysfunctions and stress.^5^ Therefore, the catalytic mechanism and the inhibition of the bc_1_ complex have important biomedical^6^ and biotechnological^7^ applications.

The structure of the bc_1_ dimer is well-established (Figure 1A).^1,8,9^ The Q substrate binds to the Q_*o*_ active site in its reduced quinol form (dihydroquinone, QH_2_; Fig. 1B). Extensive spectroscopic, kinetic, and mutational studies have identified the adjacent heme b_*L*_ in the cytochrome b unit and [2Fe-2S] cluster in the Rieske protein as electron acceptors for quinol oxidation (Fig. 1C).^9^ Electron transfer in the Q_*o*_ site is bifurcated, with each electron proceeding to a different metal center via quantum tunneling.^10–12^ Due to the probabilistic nature of this process, it is unlikely that both transfers occur simultaneously, requiring an intermediate and transient semiquinol radical (one-electron oxidized). ^9^

**Figure 1:**
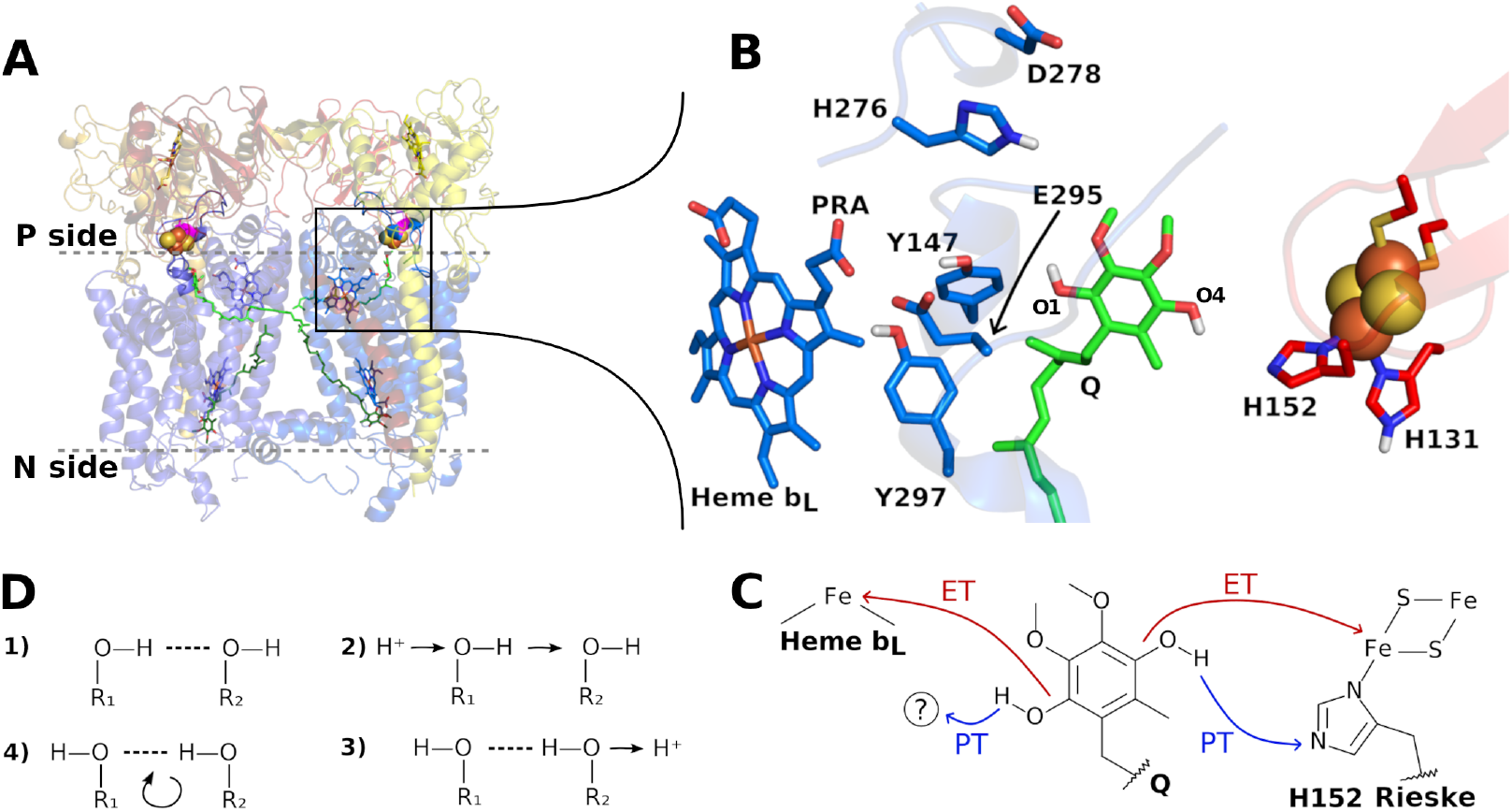
Structure of cytochrome *bc*_1_ and reactivity of its Q_*o*_ site. **A**) Essential catalytic units in the *bc*_1_ dimer from *R. sphaeroides*^8^ are shown in cartoon with cytochrome b (cyt b) in blue, cytochrome c_1_ in yellow, and Rieske protein in red. Membrane interface is in gray dashes with Q molecules in green sticks bound to the Q_*o*_ site (membrane P side) in quinol form and bound to the Q_*i*_ site (N side) in quinone form. FeS clusters are in orange and yellow spheres, hemes b_*L*_ (near site Q_*o*_) and b_*H*_ (near site Q_*i*_) in blue sticks, hemes c in yellow and cyt b D278 in magenta. **B**) Close and rotated view of reactive groups in the Q_*o*_ site with labels in residues and in phenolic oxygens of the quinol substrate (QH_2_). **C**) Electron (ET) and proton (PT) transfers for complete quinol oxidation. The question mark indicates that acceptors of one phenolic proton are unknown and investigated here. **D**) Four steps of a Grotthuss proton hopping between two hydrogen-bonded groups.

Two chemical protons are also released from quinol upon oxidation (Fig. 1C). However, the groups within the Q_*o*_ site that participate in binding and transferring these protons to the bulk solvent on the positive side of the membrane remain poorly understood.^9,13^ Identifying these groups and elucidating the proton transfer (PT) mechanism in the *bc*_1_ complex are the primary objectives of this work.

Long-range proton transfer in aqueous solutions and within solvated protein channels or cavities occurs through proton-conducting wires composed of water molecules and protonable groups connected by sequential hydrogen bonds (H-bonds, step 1 in Fig. 1D).^14^ Originally proposed by Grotthuss,^15^ the conduction of an excess H^+^ through the wire involves a series of bond-breaking and bond-forming events, or proton hops, along the wire (steps 2 and 3 in Fig. 1D), followed by the reorientation of participating groups (step 4, returning to step 1 in Fig. 1D). Thus, the composition of the proton wire and its interactions with the surrounding environment are critical for efficient proton transport.

The energetics of proton transfer along these wires can be estimated using molecular simulations with hybrid quantum chemical / molecular mechanical (QC/MM) potentials^16^ or related methods.^17^ The highly concerted Grotthuss hopping is effectively modeled using specialized reaction coordinates, such as the modified center of excess charge (mCEC),^18,19^ which capture the non-linear, three-dimensional nature of proton wires and the non-local dynamics of the transfer process. Notably, this coordinate does not require prior identification of specific atom pairs involved in each transfer step. Instead, hopping sequences and the participation of particular groups naturally emerge from the simulated reaction. Solvation dynamics and interactions with the wire environment should be reasonably sampled, so an efficient quantum chemical (QC) treatment is also required.

In previous studies, we used extensive molecular dynamics (MD) simulations to characterize interactions within the Q_*o*_ site.^11,20^ These simulations identified high flexibility among side chains, leading to three binding modes for the quinol substrate. Two of these modes appear proximal to heme b_*L*_ and enable electron and proton transfer, while a third mode represents a pre-reactive state, in agreement with recent cryoEM structures with a bound Q.^21–23^ Additionally, our MD results show that the Q_*o*_ site is highly hydrated, with several water molecules creating an H-bond network connecting quinol with conserved residues such as Y147, E295, and Y297 (residue numbering from the *bc*_1_ complex of *R. sphaeroides*, Fig. 1B). Through this network, at least five distinct proton wires were identified that could facilitate proton transport to final acceptors like H152, D278 and the heme b_*L*_ propionate-A group (PRA_*bL*_) that then release the protons to the bulk water.

Here, the energetics of proton transfer (PT) from the Q substrate in various redox states, across the multiple proton wires identified at the Q_*o*_ site, are evaluated to determine the relative stability and composition of the wires, as well as the sequence of proton hopping events. To simulate PT with greater accuracy, free energy profiles were obtained using hybrid QC/MM simulations and the mCEC reaction coordinate as mentioned above. The results confirm that heme PRA_*bL*_ can serve as a final proton acceptor via energetically favorable pathways, whereas D278 appears less likely to fulfill this role. Y147 is shown to be essential for complete PT, acting as an intermediary proton relay in all identified wires. The proton wires exhibit redundancy, functioning as a robust proton-conducting network resilient to mutations. Based on these findings, a sequence of reactions constituting the Q-cycle at the Q_*o*_ site is proposed, which may also help to elucidate the Q redox chemistry in other respiratory enzymes.

## Methods

### Set-up of molecular models

The cytochrome *bc*_1_ model used here to simulate PT reactions was based on the x-ray crystal structure of *Rhodobacter sphaeroides* (PDB 2QJP^24^) with ubiquinol-6 (with 6 isoprenoid units) bound to the Q_*o*_ site, and embedded in a solvated POPC lipid membrane. This model system was relaxed and equilibrated in previous classical MD simulations, where details of model construction were provided.^20^ Three configurations, representative of the local conformation and hydration of key residues in the Q_*o*_ site for the two reactive binding modes identified,^20^ were extracted from these trajectories at 147.0 ns from cyt b chain A (referred to as 147A), at 141.0 ns (141D) and 152.9 ns (152D) from cyt b chain D (Table S1) and treated equivalently herein for the PT simulations.

Each model was built by centering the initial geometry at the phenolic oxygen O1 of QH_2_ in the studied Q_*o*_ site. All atoms beyond r_*cut*_ = 32 Å, plus two iterations of bonded neighbors, were truncated, leaving approximately 15,800 atoms in the model. O1 in QH_2_ is positioned midway between the Rieske protein H152 and the cyt b heme PRA_*bL*_. In all simulations, atoms located further than r_*move*_ = 16 Å from O1, plus two iterations of bonded neighbors, were kept frozen, while the remaining atoms were relaxed for 5 ps of molecular dynamics with the QC/MM potential.25–27 Simulated reactant states consisted of a combination of two protonation and three redox forms: Before the first PT, the Q substrate was modeled in the double-protonated form (QH2, Fig. 2A), with the Rieske H152 N_*ϵ*_ de-protonated. After the first PT to H152 (Fig. 2B), Q was in the mono-protonated form (QH) and H152 N_*ϵ*_ was protonated. The three Q substrate redox states were as follows: fully- or double-reduced Q with oxidized FeS center and heme b_*L*_ (Q_*red*_ state); one-electron oxidized Q radical with reduced FeS center and oxidized heme b_*L*_ (Q_*semi*_); and two-electron oxidized Q with reduced FeS center and heme b_*L*_ (Q_*oxi*_).

**Figure 2:**
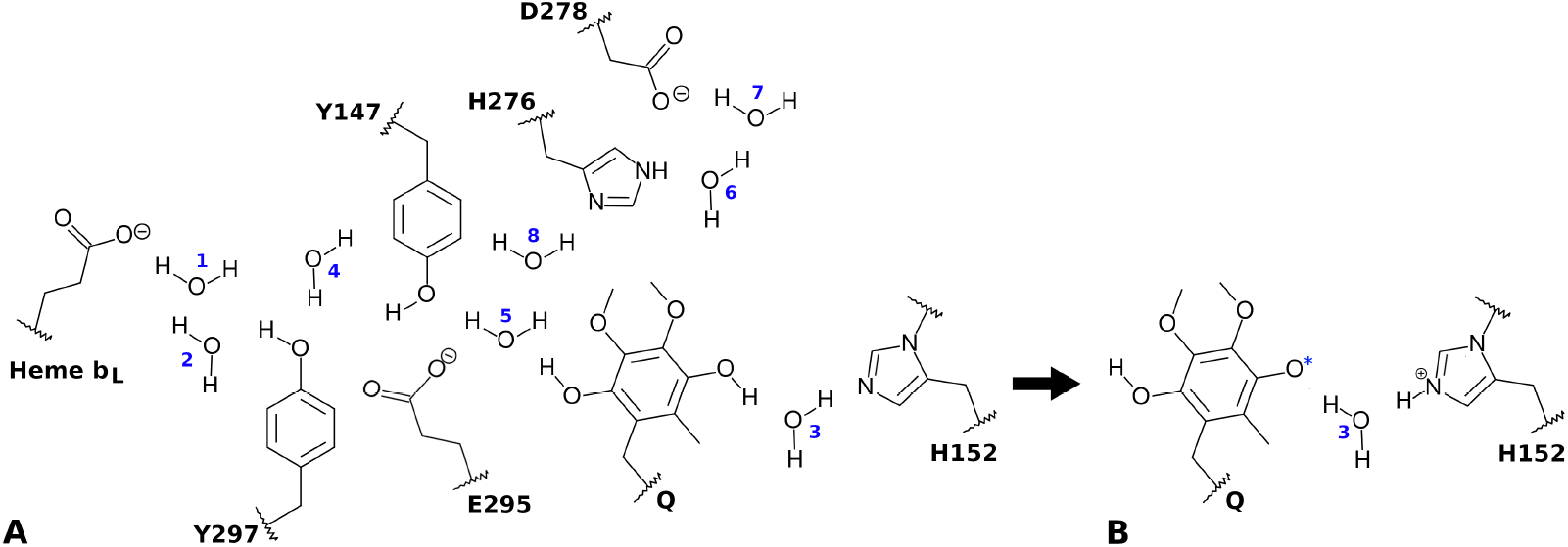
H-bond network and reactive centers in the Q_*o*_ site. For each QC/MM simulation, only groups and labeled waters given in Table 2 were represented in the QC region. **A**) Reactant state simulated in Fig. 3, corresponding to the state before the first PT. **B**) Reactant state for the second PT simulated in Figs. 4–6, after the first PT to H152 and with all other groups as shown in **A**. The asterisk in O4 indicates that various oxidation states were simulated.

### Hybrid QC/MM potential

Various QC regions were studied (Fig. 2 and Table 2). The hydrophilic Q-head of Q was always modelled in the QC region and the hydrophobic Q-tail in the MM region, with the QC/MM boundary at the C7–C8 bond (first isoprenoid group). The side chain of Rieske H152 was also always in the QC region and its N_*δ*_–Fe bond was capped with a hydrogen link-atom, with the FeS center (including both iron and sulfide atoms) and its remaining ligands modeled in the MM region. For heme b_*L*_, only the PRA_*bL*_ group was in the QC region, capped at the C_*α*_–C2 bond. All protein side chains in the QC region had their boundary placed at the C_*α*_–C_*β*_ bond. Water molecules shown in Fig. 2 were included in the QC region accordingly to their labels and Table 2. Extra water molecules were added to the QC region in comparison to the minimum number of waters previously identified for each proton wire (Table 1).^20^

**Table 1:** Proton wires previously identified^20^ for transfer between the quinol donor and various acceptors in the Q_*o*_ site, with intermediate residues and water molecules (minimum number:labels given in Fig. 2) required for proton hopping.

| Wire | Donor | Residues | Water | Acceptor |
| --- | --- | --- | --- | --- |
| A | $Q_{O4}$ | | 1: 3 | H152$N_\epsilon$ |
| B | $Q_{O1}$ | Y147, E295, | 1: 1 | PRA$_{bL,O\gamma}$ |
| C | $Q_{O1}$ | Y147, Y297 | 2: 1,5 | PRA$_{bL,O\gamma}$ |
| D | $Q_{O1}$ | Y147 | 3: 1,4,5 | PRA$_{bL,O\gamma}$ |
| E | $Q_{O1}$ | Y147, E295, H276 | 3: 5,6,7 | D278$O_\delta$ |

**Table 2:** Details of the QC region in hybrid QC/MM simulations. The Figure column shows the corresponding figure and panel in the Results. Initial is the configuration used to start simulations (Table S1). QC groups and QC water molecules (total number:labels given in Fig. 2) represented in the QC region. QC(charge, multi) gives the total charge and spin multiplicity in the QC region.

| Figure | Initial | QC groups | QC waters | QC(charge, multi) |
| --- | --- | --- | --- | --- |
| 3A | 141D | Q, H152, PRA<sub>bL</sub>, Y147, E295 | 3:1,3,5 | (−2, 1) |
| 3A | 152D | Q, H152, PRA<sub>bL</sub>, Y147 | 5:1,3,2,4,5 | (−1, 1) |
| 3B | 152D | Q, H152, PRA<sub>bL</sub>, Y147 | 5:1,3,2,4,5 | (0, 2) |
| 3B | 147A | Q, H152, PRA<sub>bL</sub>, Y147, Y297 | 3:1,3,5 | (0, 2) |
| 3C | 147A | Q, H152, PRA<sub>bL</sub>, Y147, Y297 | 3:1,3,5 | (0, 1) |
| 3C | 141D | Q, H152 | 1:3 | (0, 1) |
| 3D | 147A | Q, H152, PRA<sub>bL</sub>, Y147, Y297 | 3:1,3,5 | (0, 2) |
| 4A | 141D | Q, H152, PRA<sub>bL</sub>, Y147, E295 | 3:1,3,5 | (−1, 2) |
| 4B | 141D | Q, H152, PRA<sub>bL</sub>, Y147, E295 | 3:1,3,5 | (−2, 1), (0, 1) |
| 4C | 152D | Q, H152, PRA<sub>bL</sub>, Y147 | 5:1,3,2,4,5 | (0, 2) |
| 4D | 152D | Q, H152, PRA<sub>bL</sub>, Y147 | 5:1,3,2,4,5 | (−1, 1), (+1, 1) |
| 4E | 147A | Q, H152, PRA<sub>bL</sub>, Y147, Y297 | 3:1,3,5 | (0, 2) |
| 4F | 147A | Q, H152, PRA<sub>bL</sub>, Y147, Y297 | 3:1,3,5 | (−1, 1), (+1, 1) |
| 5A | 141D | Q, H152, PRA<sub>bL</sub>, E295 | 3:1,3,5 | (0, 1) |
| 5B | 152D | Q, H152, PRA<sub>bL</sub>, | 5:1,3,2,4,5 | (0, 2) |
| 5C | 152D | Q, H152, PRA<sub>bL</sub>, | 5:1,3,2,4,5 | (+1, 1) |
| 6A | 141D | Q, H152, Y147, E295, H276, D278 | 5:3,5,6,7,8 | (−1, 2) |
| 6B | 141D | Q, H152, Y147, E295, H276, D278 | 5:3,5,6,7,8 | (0, 1) |

The efficient PM6 semi-empirical method^28^ was used for the QC region. Comparison with higher levels of theory (DFT) suggests that this treatment is reasonably accurate (Fig. S1). For the MM region and Lennard-Jones parameters of QC centers in QC/MM simulations, the same force-field was applied as in previous classical MD simulations,^20,29,30^ using CHARMM36m^31^ and TIP3P^32^ parameters for protein, lipids and water, with recalibrated parameters for FeS ligands and heme b centers.^20^ Hydrogen link-atoms were used to cap valencies in the QC region.

For simulations with Q in the partially oxidized semiquinol form (Q_*semi*_), it was assumed that electron transfer proceeded to the FeS center, reducing the partial charge of FE2 (Cys-bound) by −0.50 |e| and each inorganic sulfide by −0.25 |e|. For simulations with Q fully oxidized (Q_*oxi*_), it was assumed both the heme b_*L*_ and the FeS center were reduced, reducing the partial charge of heme FE by −0.60|e| and each bound nitrogen in the porphyrin ring by −0.10 |e|. These changes were determined after comparing partial charges in oxidized and reduced states from DFT calculations on model compounds for FeS and heme centers and were assigned before any QC/MM boundary charge redistribution.^33^ Given that these centers are spatially distant and separated by multiple covalent bonds from the reactive protons, any inaccuracies in their MM parameters are expected to have minimal impact on the simulation results.

Standard electrostatic QC/MM embedding was used, with long-range electrostatics cut off by an atom-based force-switching function (r_*on*_ = 8 Å and r_*off*_ = 12 Å). This cut-off aligns with the model setup, ensuring that all moving atoms remain within the switching range of long-range interactions, r_*off*_ < (r_*cut*_ − r_*move*_). To prevent diffusion into the MM region, a flat-bottom harmonic potential 2.0 Å wide with a 150 kJ mol^*−*1^ Å^*−*2^ force constant was applied to the oxygens of all QC waters, centered at their initial positions. All QC/MM simulations and analysis were conducted with the pDynamo library version 3, ^33,34^ where the mCEC^18^ reaction coordinate was implemented (see below). Initial configurations and sample pDynamo scripts with all simulation parameters are available online^35^ to enable full reproduction of this study.

### Umbrella sampling and reaction coordinate

Proton transfer reactions were simulated with these models and QC/MM potential by umbrella sampling (US) ^36^ with Langevin MD, temperature of 310 K, collision frequency of 25.0 ps^*−*1^ and a time-step of 1 fs. Umbrella sampling was performed along the ζ reaction coordinate defined below (eq. 2). The ζ=[0,1] interval was divided in equally spaced windows of 0.1 length and a force constant k_*umb*_ = 1000 kJ mol^*−*1^ Å^*−*2^ was employed. If necessary, additional windows were included to increase ζ overlap with k_*u*_*′* _*mb*_ = 3k_*umb*_. Each window was sampled by at least 0.5 ns, which is shown in Fig. S2 to be enough for convergence of the profiles. Some simulations had sampling increased to 1 ns per window as indicated. Free energy profiles were pieced together using the weighted histogram analysis method (WHAM)^37^ with the initial 0.1 ns of each US window discarded for equilibration. Statistical uncertainties were estimated as 95% confidence intervals by bootstrap analysis with 30 resampling cycles. ^38^

The modified center of excess charge (mCEC)^18,19^ was implemented in the pDynamo 3 library and used here as a reaction coordinate to drive global and non-linear proton-transfer in Grotthuss proton hopping (Fig. 1D) through the identified wires. Briefly, the mCEC is defined as:

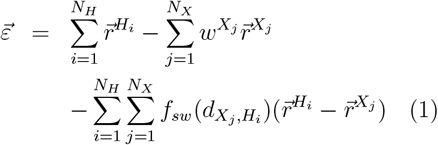

where *X*_*j*_ represents the atom bound to a ionizable proton (H_*i*_), 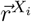 is the position of X_*i*_ and w^*Xj*^ is the weight associated with atom *X*_*j*_. This is defined by the least number of protons bound to that atom. f_*sw*_ is a smooth switching function depending on A to B atom distance (d_*A,B*_). Here, values of r_*sw*_ = 1.20 Å and d_*sw*_ = 0.04 Å were used in the f_*sw*_ function.^18^ Weights were assigned as 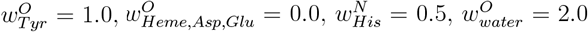 and 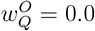.

As the center of charge 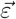 corresponds to a spatial (vector) coordinate, we employ the relative coordinate ζ to enhance sampling in US simulations:

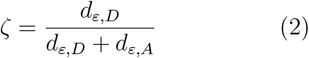

where d_*ε,D*_ and d_*ε,A*_ are distances between 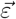 and the position of the proton donor or the acceptor atoms, respectively. ζ is defined in the interval [0,1], with 0 corresponding to the reactant state (pre-proton transfer) and 1 is the product state (post-proton transfer). In QC/MM simulations performed here, this type of reaction coordinate described transfer for different donor (D) and acceptor (A) pairs: from donor Q_*O*1_ to acceptor heme PRA_*bL,Oγ*_ (ζ coordinate) in Figs. 3A,B, 4 and 5; from Q_*O*1_ to D278_*Oδ*_ (ζ^*′*^ coordinate) in Fig. 6; and from Q_*O*4_ to H152_*Nϵ*_ (ζ^*′′*^ coordinate) in Fig. 3C,D. All QC waters and protonable groups listed in Table 2 were included in the respective definition of mCEC coordinate.

**Figure 3:**
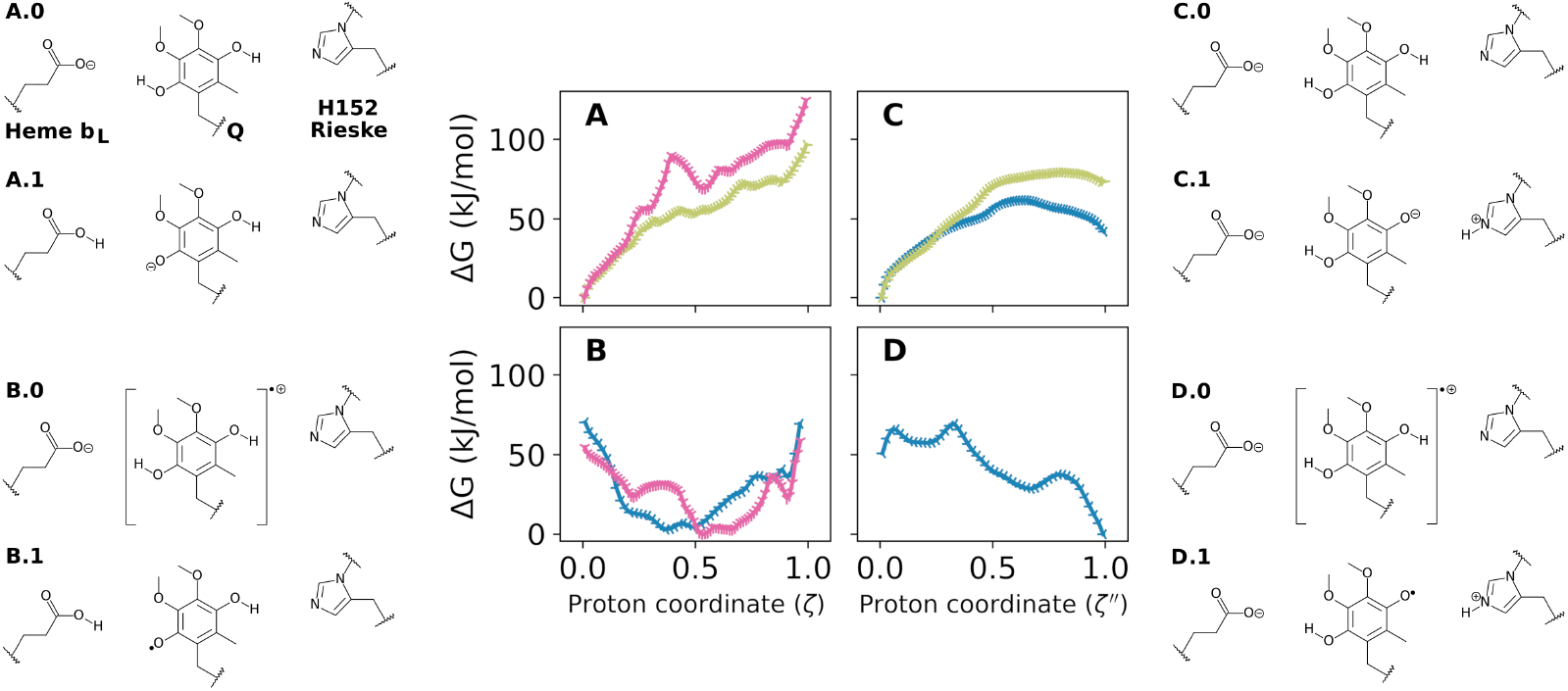
Reactivity of the first proton transfer from the quinol substrate depicted as free energy profiles. PT was simulated from double-protonated and: **A**) double-reduced quinol (QH_2_) to PRA_*bL*_; **B**) semiquinol radical 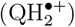 to PRA_*bL*_; **C**) double-reduced quinol (QH_2_) to H152; and **D**) semiquinol radical 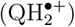 to H152. Charge states of PRA_*bL*_, Q and H152 groups in the reactant (proton coordinate ζ = 0→X.0) and product (ζ = 1→X.1) are shown besides the respective profile X. Simulations were performed with hybrid QC/MM potentials with QC region defined in Table 2 and Fig. 2A. Profile colors denote different wires and initial configurations with 147A in blue, 141D in gold and 152D in pink. Statistical uncertainties in the free energies are approximately ± 1 kJ/mol and are smaller than the symbols shown in each panel.

## Results

A set of 21 free energy profiles of proton transfer reactions in the Q_*o*_ site is presented here probing various proton wires (Table 1)^20^ and reactant states (Tables 2 and S1) for quinol oxidation. First, the acceptor group for the initial PT is identified, along with its product which serves as a reactant state for the subsequent second PT. Then, proton wires and acceptors for the second PT are tested to define the final secondary acceptor, the reactive composition of wires, and the expected proton hopping sequence. Detailed reaction mechanisms, describing bond breaking and formation sequences, will be presented for a select subset of reactions, particularly those with favorable energetics (*e*.*g*., Fig. 4A) and thus more likely to occur.

**Figure 4:**
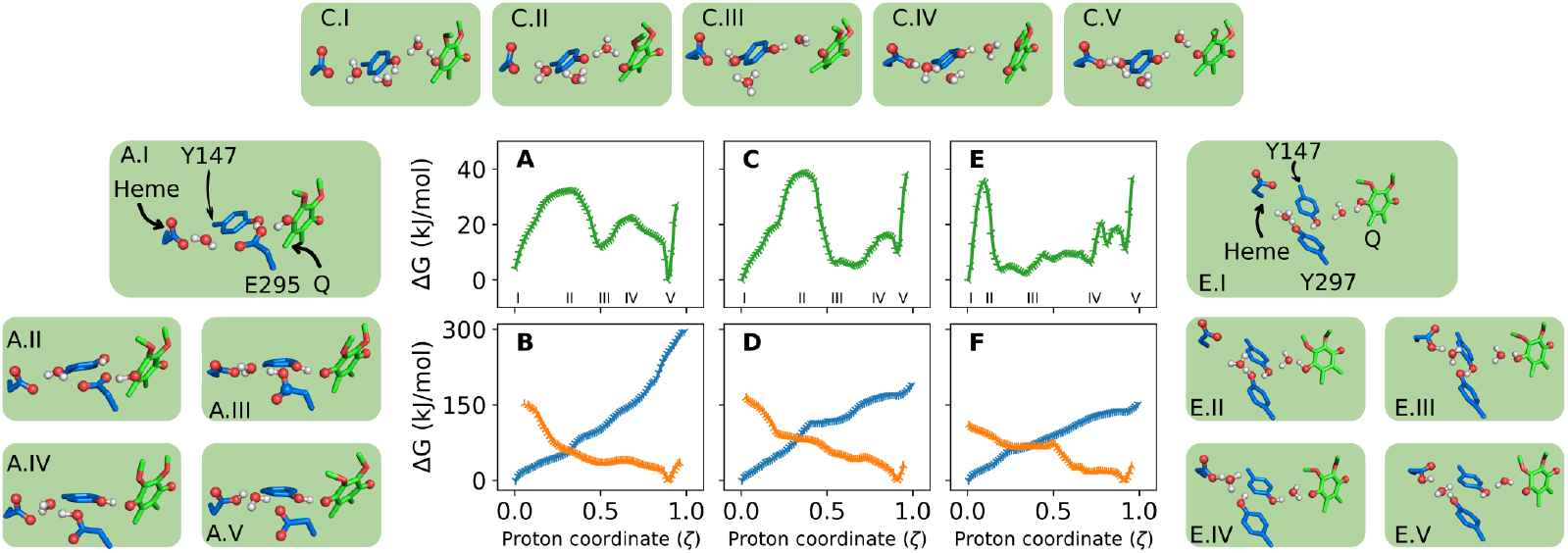
Reactivity of the second proton transfer to heme A-propionate (PRA_*bL*_) depicted as free energy profiles. Three different proton-wires from mono-protonated QH (Fig. 2B) were simulated via: **A**,**B**) Y147 and E295 (wire B in Table 1); **C**,**D**) Y147 only (wire D); and **E**,**F**) Y147 and Y297 (wire C). Colors denote the redox state of QH, with semiquinol (QH^*•*^) in green, reduced (QH^*−*^) in blue and fully oxidized (QH^+^) in orange. Boxed inserts show representative structures for reactive groups at proton coordinates indicated by roman numerals in the respective plot X-axis. For example, inset A.III corresponds to the intermediate found at ζ=0.5 in panel **A**.

### First proton transfer proceeds to His152 and generates a mono-protonated semiquinol radical

The initial PT from the fully protonated quinol substrate was simulated and the estimated free energy profiles are shown in Fig. 3. Using fully reduced QH_2_ as the donor and heme PRA_*bL*_ as the acceptor, Fig. 3A shows steep uphill profiles with high free energies (over 100 kJ/mol) and no barrier for reverse PT, revealing an unstable product. This indicates that PT is unlikely for this reactant state and proton wires. Profiles are qualitatively similar for the two simulations, supporting that results are consistent across wires with different composition and initial configuration.

Fig. 3B shows profiles for the same donor-acceptor pair, but now starting from a semiquinol radical reactant 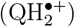, with one-electron oxidized by the high-potential Rieske FeS center. A pronounced free energy minimum (∼50 kJ/mol) is observed at proton coordinate ζ ∼0.5, corresponding to the transferred charge being trapped as a hydronium (H_3_O^+^) H-bonded to Y147 and the PT reaction does not complete to heme PRA_*bL*_. Again, profiles are qualitatively similar for PT through two different wires and initial configurations.

Fig. 3C,D are the only plots in this study describing PT to Rieske H152, from donor Q_*O*4_ to acceptor H152_*Nϵ*_ connected by a water molecule in bridge, simultaneously H-bonded to both centers (Fig. 2A and proton wire A in Table 1). The profiles in Fig. 3C for PT from QH_2_, thus before any ET from the substrate, show a high barrier (>60 kJ/mol) and an unstable product, suggesting again that PT should not be observed before quinol oxidation.

Fig. 3D was obtained for PT from 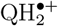 to H152, after one-electron oxidation of the quinol substrate by the FeS center. The profile shows a small barrier (<20 kJ/mol) and high stability (<−50 kJ/mol) for PT, suggesting that reaction would be fast and stable. The simulated reaction mechanism for this PT is simply a proton donation from Q_*O*4_ to the bridge water oxygen, which in concert donates another proton to H152_*Nϵ*_, similar to a water asymmetric bond-stretch. The transition state at ζ^*′′*^ = 0.35 corresponds to a transient hydronium ion formed along this concerted transfer.

Thus, the only feasible and efficient reaction identified for the initial PT from quinol involves transfer from the one-electron oxidized 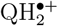 to protonate H152_*Nϵ*_. This reaction produces a mono-protonated semiquinol radical QH^*•*^.

### Second proton transfer proceeds to the propionate in heme *b*_*L*_ by three different wires

All simulations for the second PT in the Q_*o*_ site begin from a reactant state with the mono-protonated QH species (proton bound to O1, with O4 ionized), as shown in Fig 2B. The free energy profiles in Fig. 4 describe PT to the heme PRA_*bL*_ through three different proton wires (labeled B, C, and D in Table 1), involving E295, Y297, and Y147 side chains and nearby water molecules.^20^ All possible QH redox states are tested and the behavior for each state is qualitatively similar across the three wires.

If the reactant is fully reduced (QH^*−*^, blue profiles), the PT reaction free energies exceed 150 kJ/mol, indicating that the second PT would not occur, consistent with results from Fig. 3A,C for the first PT. This outcome strongly indicates that proton transfer from Q is unlikely to proceed in the Q_*o*_ site *without* prior or concerted electron transfer.

In the double-oxidized reactant (QH^+^, orange profiles), the free energy for PT is barrier-less and downhill, leading to a stable, protonated heme b_*L*_ product (ζ=0.9). However, the released “hot-proton” would transfer to any protonable group, making it highly improbable for the native Q-cycle to rely on such an irreversible and highly dissipative (exergonic) reaction.

Conversely, in the one-electron oxidized form (QH^*•*^, green profiles), low barriers (25-40 kJ/mol) emerge at early proton coordinates, leading to a protonated heme PRA_*bL*_ product, in a nearly equilibrium or ther-moneutral reaction (ΔG_*R*_ =± 10 kJ/mol). This suggests a conservation of free energy and a reversible transformation, ^39,40^ supporting a much more plausible pathway for the Q-cycle.

For the proton wire B via Y147 and E295 (Fig 4A and A.I-A.V insets), PT proceeds from QH^*•*^ with an early barrier (A.II) corresponding to breaking the O4–H bond and forming an intermediate protonated at the E295 side chain (A.III). The proton bound to E295 may come either directly from Q_*O*4_ or passed concertedly to Y147_*OH*_ which transfer the excess proton to E295. This intermediate is de-protonated by a bridging water passing the excess proton to heme (A.V).

In wire D, the role of E295 as an intermediate proton acceptor is substituted by water molecules (Fig 4C and C.I-C.V insets) that help to bridge PT from QH^*•*^ to Y147 (C.II), form a hydronium intermediate (C.III) instead of a protonated E295 and transfer the excess charge to heme (C.IV).

For the proton wire C via Y147 and Y297 (Fig 4E and E.I-E.V insets), the early barrier corresponds to formation of an extended H-bond network involving O4_*Q*_, the two Tyr_*OH*_ and the oxygens of two water molecules (E.II). An intermediate with one proton shared between Y147 and Y297 is formed with heme PRA_*bL*_ already protonated (E.III). Complete PT is achieved after rearrangement of the H-bond pattern (E.IV), reminiscent of a proton-hole transfer.^41^

### Residue Y147 is necessary to complete the second proton transfer

To investigate the chemical role of Y147 in the second PT reaction, simulations shown in Fig. 5 were conducted using the same proton wires (B and D, Table 1), but with an unreactive Y147 now modeled in the MM region (Table 2). This mimics the effect of a Y147 point mutation to a residue unable to exchange protons, but with minimal disruption to the surrounding chemical environment. Y147 in the MM region can not form covalent bonds but can still form H-bonds and interact electrostatically with groups in the Q_*o*_ site.

**Figure 5:**
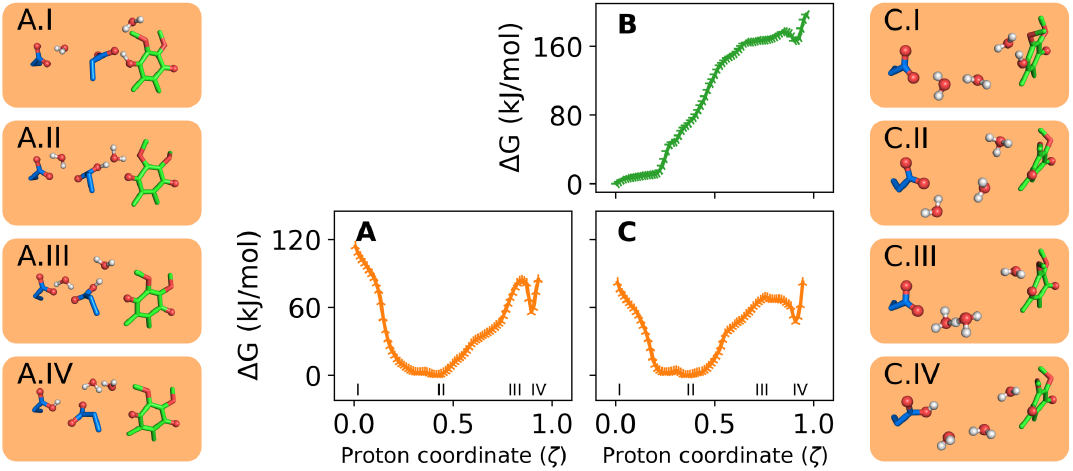
Reactivity of the second proton transfer to PRA_*bL*_ without an intervening Y147. Two proton-wires were simulated: **A**) PT via E295 (wire B without Y147); and **B**,**C**) PT via water only (wire D without Y147). Colors denote the QH redox state with semiquinol (QH^*•*^) in green and fully oxidized (QH^+^) in orange.

The free energy profile in Fig. 5A shows a striking difference compared to that in Fig. 4B for the proton wire B. The initial PT from oxidized QH^+^ is again highly exergonic, forming a protonated E295 (inset A.II). However, the PT reaction does not complete to heme PRA_*bL*_ due to a substantial barrier (95 kJ/mol, A.III) and the instability of the protonated PRA_*bL*_ (A.IV). As a result, the excess proton becomes trapped in E295 when Y147 is chemically inactive. A similar pattern occurs in Fig. 5C (wire D as in Fig. 4D), but here trapping involves a hydronium ion (inset C.II). For reaction of the singly oxidized QH^*•*^ (Fig. 5B), the free energies for PT to heme are over 160 kJ/mol, preventing any reaction. This outcome contrasts sharply with the PT behavior through the same wire when Y147 is chemically active (Fig. 4C).

### Transfer to D278 is not energetically feasible

A recent study proposed that H276 and D278 in cyt b may function as the proton acceptors for the second PT.^42^ Hydration of the Q_*o*_ site revealed a network of H-bonds connecting the Q substrate to H276 and D278 in cyt b via 2 or 3 water molecules (Fig. 1B and wire E in Table 1),^20^ supporting a Grotthuss proton hopping mechanism.

The energetics of this proton wire E is shown in Fig. 6. For the semiquinol QH^*•*^ reactant, transfer proceeds from Q_*O*4_ by a small barrier (10 kJ/mol) via an Eigen-type ion (A.II), leading to a stable intermediate with the H276 side chain double-protonated (A.III). Proton transfer to D278 occurs through a water bridge but requires a high free energy (80 kJ/mol) and results in an unstable product (A.IV). For the double oxidized QH^+^ reactant, an analogous intermediate forms after a downhill transfer, with the excess proton again becoming trapped in H276.

**Figure 6:**
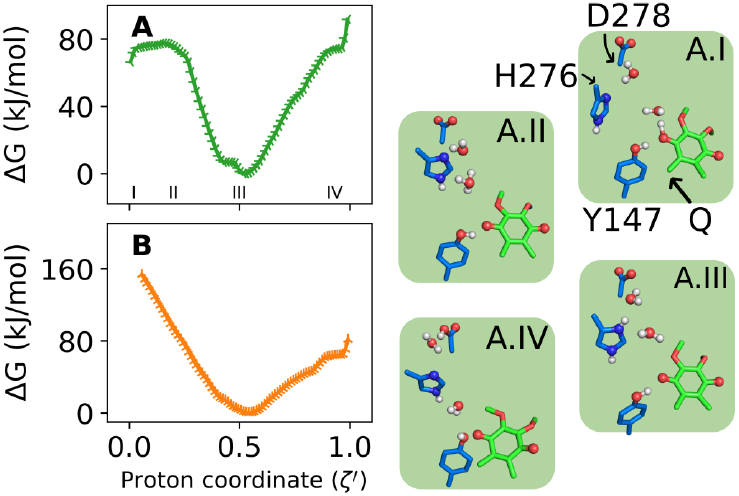
Reactivity of the second proton transfer to D278. One proton wire (E, Table 1) was simulated. Colors denote the QH redox state with semiquinol (QH^*•*^) in green and fully oxidized (QH^+^) in orange.

## Discussion

A comprehensive set of reactant states, proton wires, and simulation set-ups for possible PT reactions in the Q_*o*_ site of cytochrome *bc*_1_ was investigated by hybrid QC/MM simulations and a specialized reaction coordinate (mCEC).

The statistical uncertainties for the calculated free energies are approximately ±1 kJ/mol. This represents a lower bound of the total uncertainty, which is challenging to assess due to the approximate nature of the simulations. The QC level employed in the QC/MM potential (see Methods) is one possible source of error. However, Fig. S1 shows that this treatment is reasonably accurate compared to higher levels of theory (DFT), indicating a minor contribution to overall uncertainty.

Another possible concern is the use of configurations sampled in the QH_2_ reactant state (Table S1) as starting structures for the simulations of the second PT (Figs. 4-6). However, this should not be a limitation, as the first ET and PT reactions are rapid (nanosecond time scale or faster) and no significant structural rearrangements are likely to occur within such a short period. Furthermore, the initial 100 ps of each US simulation window was discarded before estimating the free energy profiles, allowing sufficient time for relaxation and equilibration of the Q-head, side chains and water molecules in response to changes in the substrate reactant state.

While other sources of error may still be present (*e*.*g*., in the molecular model set-up), the results presented here remain consistent across different models and initial configurations, providing strong qualitative reliability. Therefore, all discussions and conclusions will emphasize qualitative and relative trends, such as uphill vs. downhill profiles, small vs. large barriers, and complete vs. trapped reactions.

Analysis of free energy profiles presented below (after Fig. 7) is based on the following (bio)energetic rationale: highly uphill profiles (> 70 kJ/mol) imply that the process is energetically unfeasible. Low barriers (< 15 kJ/mol) for the reverse reaction indicate an unstable product. Trapped intermediates suggest that the transfer may remain incomplete, allowing the accumulation of intermediates, with possible reversibility or short-circuiting of the reaction. In these three cases, the simulated process or step is unlikely to occur or reach completion under physiological operation of the Q-cycle.

**Figure 7:**
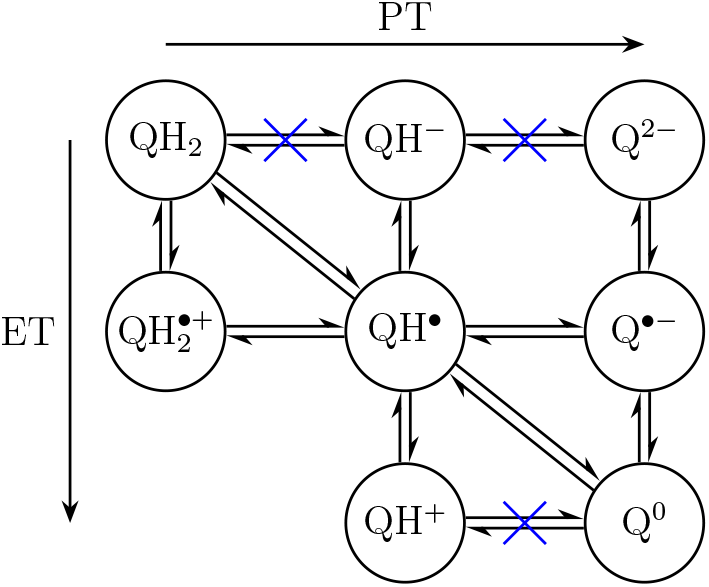
Possible reactions for quinol oxidation. Horizontal equilibria represent proton transfer (PT) reactions, which were investigated here. Vertical equilibria denote electron transfer (ET) and diagonal indicate coupled proton-electron transfer (CPET). Equilibria marked with a blue cross are unlikely to occur in the Q_*o*_ site under normal operation of the Q-cycle based on the analysis presented here.

Moreover, steep downhill profiles suggest that a transfer would occur indiscriminately, with a “hot-proton” transferred to any acceptor group. These profiles dissipate substantial free energy, often exceeding what is provided by complete QH_2_ oxidation in the *bc*_1_ complex (∼70 kJ/mol)^9^ and again, the respective transfer step is unlikely to occur physiologically.

Plausible PT steps for the Q-cycle, however, should exhibit profiles with low to medium barriers (up to ∼ 45 kJ/mol) and operate near equilibrium or thermoneutral conditions (reaction |ΔG_*R*_|< 20 kJ/mol), with both conditions also required for the Q-cycle to operate in reverse over the same steps.

Fig. 7 shows possible ET and PT reactions for complete quinol oxidation (or quinone Q^0^ reduction in reverse). Horizontal equilibria describe PT reactions, which were all tested here in the Q_*o*_ site for different proton acceptors (H152, PRA_*bL*_ and D278).

For the first PT (left to middle column in Fig. 7), the reaction does not proceed to heme PRA_*bL*_ or to H152 before one-electron oxidation of quinol by the FeS center (Fig. 3). After the first ET, PT still does not proceed to PRA_*bL*_ but transfer to H152 is fast and thermodynamically favorable.

Previous computational studies have shown that a coupled proton-electron transfer (CPET) from quinol to H152 should also be efficient.^11,12^ These studies included a FeS center bound to H152 within the QC region and also yielded a stable semiquinol radical (QH^*•*^) as the product.

Therefore, either a stepwise (in the ET+PT sequence, but not in PT+ET) or a coupled (CPET) transfer from QH_2_ to protonate H152 are possible and yield a monoprotonated semiquinol radical (QH^*•*^) for the first oxidation step (Fig. 7). The second PT from Q_*O*1_ will proceed preferably once Q_*O*4_ is ionized and the substrate has undergone (one-electron) oxidation. These results are in line with previous experimental spectroscopic^9,43,44^ and mutational^45^ studies that suggested H152 is one of the proton acceptors. Participation of heme PRA_*bL*_ as a proton acceptor was recently proposed,^20^ based on MD simulations of local hydration in the Q_*o*_ site and the formation of an H-bond network connecting QH_2_ and PRA_*bL*_, also supported by cryoEM structures. ^21–23^ Involvement of propionate groups from redox-active hemes in PT has been suggested in other bioenergetic enzymes, including respiratory complex IV (CcO).^46–48^ However, obtaining experimental evidence for propionate participation in *bc*_1_ is challenging, as mutations affecting heme groups often render the enzyme non-functional, and may even prevent correct protein folding. ^49^

One goal of this study was to determine whether the energetics of PT to PRA_*bL*_ is consistent with the proposed role of this group as an acceptor. Indeed, simulations of the second PT to heme PRA_*bL*_ through three different proton wires confirm this (Fig. 4A,C,E, corresponding to QH^*•*^ **⇌** Q^*•−*^ in Fig. 7). Barriers between 25-40 kJ/mol and reaction free energies of ±10 kJ/mol were obtained for all three wires, indicating that this is a thermodynamically feasible and kinetically fast PT pathway. This supports the viability of heme PRA_*bL*_ as an acceptor for the second proton in the Q_*o*_ site.

These barrier heights also suggest that the lifetime of the QH^*•*^ radical intermediate is on the order of tens of nanoseconds. Thus, experimental detection of this intermediate under equilibrium conditions, which has been debated extensively in the literature,^9^ would require ultrafast spectroscopic techniques capable of capturing species within this timescale.

Acidity of QH^*•*^ in the Q_*o*_ site is markedly higher than that of QH_2_, similar to what was found from solution measurements.^50^ Notably, the nearly thermoneutral reaction energy of the QH^*•*^ **⇌** Q^*•−*^ equilibria suggests that both species will be present in the Q_*o*_ site, with the lifetime of the de-protonated radical depending on its ET rate. The thermoneutral reaction further indicate that this PT step can proceed in reverse, consistent with observations from *in vitro* experiments on reconstituted^39^ and cofactor knockout^40^ *bc*_1_ complexes.

A novel and interesting finding for the *bc*_1_ enzyme is that the three wires (B, C & D) exhibit similar energetics and could potentially function with redundancy, forming a *proton conducting network*. This is also consistent with single-point mutations in residues E295 or Y297, that still yield functional *bc*_1_ complexes.^45^ Thus, flexibility within the Q_*o*_ site, leading to variations in side chain conformations and two reactive binding modes of the Q-head,^20^ along with the operation of multiple proton wires, could all contribute a significant entropic component that enhances PT efficiency in the Q_*o*_ site.

The three proton wires (B, C & D) share two common features: the involvement of Y147 as an initial and transient proton acceptor and the delivery of the excess charge to heme PRA_*bL*_ via a water molecule. If Y147 is not chemically involved, neither PT from semiquinol QH^*•*^ or from fully oxidized QH^+^ results in complete transfer to heme PRA_*bL*_ (Fig. 5). Thus, the role of Y147 as a transient proton relay (as observed in models A.II, C.II, and E.III of Fig. 4) is critical for enabling an energetically favorable and complete PT to heme b_*L*_. Single-point mutations in Y147 have been shown to abolish or significantly impair oxidase activity in the Q_*o*_ site.^51^

A recent mutational study proposed^42^ that one of the chemical protons from quinol oxidation is transferred by H276 to D278 in cyt b. These residues are part of the previously identified H-bonded network within the Q_*o*_ site and the proton wire E (Table 1).^20^ However, PT from QH^*•*^ or QH^+^ does not proceed completely to D278; instead, the excess charge becomes trapped at H276 (Fig. 6). H276 is also unlikely to act as the proton release group due to the high free energy barrier for its deprotonation to water and its relatively distant position from bulk water. Additionally, both H276 and D278 show low conservation among *bc*_1_ variants and are frequently mutated.^20^ Therefore, it can be concluded that H276 and D278 are unlike to function as acceptors for the second chemical proton released during quinol oxidation.

For the forward quinol oxidation in the Q_*o*_ site, the results presented here (Figs. 3A,C and 4B,D,F), when analyzed using the above bioenergetic rationale, indicate that the equilibria marked by a blue cross in Fig. 7 are unlikely to occur under physiological conditions. The reverse PT from H152 and QH^*•*^ to reform QH^+^ is energetically unfavorable and kinetically slow. Furthermore, if the same reaction steps obtained for the forward cycle are mirrored in the reverse Q-cycle operation, the coupled CPET (diagonal) reaction likely represents the most probable equilibrium between the left and middle columns in Fig. 7. 11,12,44

This analysis indicates that only neutral closed-shell Q species (first and third rows in Fig. 7) are formed during both forward and reverse oxidation in the Q_*o*_ site. Double oxidation of a doubly protonated species (resulting in QH^2+^) is highly improbable and was not considered here. Protonation reactions involving QH^+^, QH^*−*^ and Q^2*−*^ are highly uphill or excessively dissipate free energy, depending on the direction considered.

Therefore, the most plausible reaction sequence for quinol oxidation in the Q_*o*_ site is QH_2_→ QH^*•*^→ Q^0^, occurring either via coupled CPET (diagonals in Fig. 7) or through a stepwise transfer involving 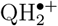 and 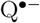 in-termediates. The coupled CPET mechanism offers the added advantage of bypassing these charged radical species and avoiding trapped intermediates (Fig. 3C).

It can be hypothesized that Q redox chemistry catalyzed by other enzymes follows a similar mechanism. In particular, respiratory complexes I and II also contain Tyr and His side chains that coordinate the Q-head within their respective Q-binding sites. ^52–54^ Based on the free energy profiles and analysis presented here, charged closed-shell intermediates such as Q^2*−*^, QH^*−*^, or QH^+^ are also unlikely to be stable within these other respiratory complexes.

Although not directly simulated in this study, it is plausible to speculate that the protons bound to H152 and PRA_*bL*_ are subsequently transferred to bulk water on the positive side of the membrane. H152 resides in the head of the Rieske domain, which is flexible and capable of moving to approach the cytochrome c_1_ heme center.^9^ This movement may expose H152 to bulk water, potentially facilitating proton release after oxidation of the FeS center.^43^ PRA_*bL*_ is located in a hydrated region directly in contact with bulk water.^20^ Consequently, the excess proton bound to this propionate can be readily released^48,55^ after heme b_*L*_ is oxidized by heme b_*H*_ (Fig. 1A).^9^ These processes would reset the redox and protonation states of the metal cofactors and protonable groups in the Q_*o*_ site, enabling another catalytic cycle following the exchange of the produced quinone (Q^0^) for a new quinol substrate.

## Conclusions

The simulations presented here integrate hybrid QC/MM potentials, a global non-linear reaction coordinate and free energy profiles to probe the energetics of possible PT reactions after quinol oxidation in the Q_*o*_ site of cytochrome *bc*_1_. Beyond classical MD simulations that revealed local hydration and possible proton wires in the Q_*o*_ site,^20^ the current qualitative analysis may validate PT pathways and exclude trapped intermediates or highly dissipative reactions.

Key reactive groups and mechanisms of proton transfer were identified for Q redox reactions. The first PT is energetically feasible to Rieske H152 after one-electron oxidation of quinol, resulting in the formation of a monoprotonated semiquinol radical (QH^*•*^). The second PT from this radical intermediate to heme PRA_*bL*_ is thermodynamic and kinetic viable, supporting the role of heme b_*L*_ as a final PT acceptor.

Residue Y147 in cyt b is a necessary initial and transient proton acceptor, without which complete PT to intermediate (such as E295) and final (heme PRA_*bL*_) acceptors is hindered. Conversely, residues H276 and D278 were found unlikely to contribute as proton acceptors.

Simulations reveal that multiple and redundant proton wires facilitate the PT reactions, creating a proton conducting network within the Q_*o*_ site, that is also robust to single-point mutations. Combined with the previously identified flexibility of side chains and reactive binding modes of the Q-head,^20^ the network of PT pathways has a significant entropic contribution to the PT efficiency in the Q_*o*_ site.

The overall mechanism suggests that the quinol oxidation sequence proceeds via QH_2_→QH^*•*^→Q^0^, without charged closed-shell intermediates. This pathway avoids high-energy charged species and reinforces an energetically optimized process for concerted proton-electron transfer in the Q_*o*_ site. These findings may be applicable to other enzymes with similar active sites and Q redox chemistry, and may help to understand proton transfer reactions in quinol-quinone redox cycling.

## Acknowledgement

Funding from Fundaçao de Amparo á Pesquisa do Estado de São Paulo (FAPESP, grant 2023/00934-5) and Conselho Nacional de Desenvolvimento Científico e TecnolÓgico (CNPq) are acknowledged.

## References

(1) Cramer, W.; Hasan, S. S.; Yamashita, E. The Q Cycle of Cytochrome bc Complexes: A Structure Perspective. Biochim. Biophys. Acta 2011, 1807, 788–802.

(2) Mitchell, P. The Protonmotive Q-cycle: A General Formulation. FEBS Lett. 1975, 59, 137–139.

(3) Osyczka, A.; Moser, C. C.; Dutton, P. L. Fixing the Q-Cycle. Trends Biochem. Sci. 2005, 30, 176–182.

(4) Crofts, A. R. The modified Q-cycle: A look back at its development and forward to a functional model. Biochim. Biophys. Acta 2021, 1862, 148417.

(5) Hamanaka, R. B.; Chandel, N. S. Mitochondrial reactive oxygen species regulate cellular signaling and dictate biological outcomes. Trends Biochem. Sci. 2010, 35, 505–513.

(6) Birth, D.; Kao, W.-C.; Hunte, C. Structural analysis of atovaquone-inhibited cytochrome bc1 complex reveals the molecular basis of antimalarial drug action. Nat. Commun. 2014, 5, 4029.

(7) Fisher, N.; Meunier, B.; Biagini, G. A. The cytochrome bc1 complex as an antipathogenic target. FEBS Lett. 2020, 594, 2935–2952.

(8) Xia, D.; Esser, L.; Tang, W.-K.; Zhou, F.; Zhou, Y.; Yu, L.; Yu, C.-A. Structural Analysis of Cytochrome bc1 Complexes: Implications to the Mechanism of Function. Biochim. Biophys. Acta 2013, 1827, 1278–1294.

(9) Sarewicz, M.; Pintscher, S.; Pietras, R.; Borek, A.; Bujnowicz, L., Hanke, G.; Cramer, W. A.; Finazzi, G.; Osyczka, A. Catalytic Reactions and Energy Conservation in the Cytochrome bc1 and b6f Complexes of Energy-Transducing Membranes. Chem. Rev. 2021, 121, 2020–2108.

(10) Hagras, M. A.; Hayashi, T.; Stuchebrukhov, A. A. Quantum Calculations of Electron Tunneling in Respiratory Complex III. J. Phys. Chem. B 2015, 119, 14637–14651.

(11) Camilo, S. R. G.; Curtolo, F.; Galassi, V. V.; Arantes, G. M. Tunneling and Nonadiabatic Effects on a Proton-Coupled Electron Transfer Model for the Qo Site in Cytochrome bc1. J. Chem. Inf. Model. 2021, 61, 1840–1849.

(12) Barragan, A. M.; Soudackov, A. V.; Luthey-Schulten, Z.; Hammes-Schiffer, S.; Schulten, K.; Solov’yov, I. A. Theoretical Description of the Primary Proton-Coupled Electron Transfer Reaction in the Cytochrome bc1 Complex. J. Am. Chem. Soc. 2021, 143, 715–723.

(13) Springett, R. The Proton Pumping Mechanism of the bc Complex. Biochim. Biophys. Acta 2021, 1862, 148352.

(14) Silverstein, T. P. The Proton in Biochemistry: Impacts on Bioenergetics, Biophysical Chemistry, and Bioorganic Chemistry. Front. Mol. Biosci. 2021, 8, 764099.

(15) de Grotthuss, C. J. T. Sur la décomposition de l’eau et des corps qu’elle tient en dissolution á l’aide de l’électricité galvanique. Ann. Chim. (Paris) 1806, 58, 54–73.

(16) Kubar, T.; Elstner, M.; Cui, Q. Hybrid Quantum Mechanical/Molecular Mechanical Methods For Studying Energy Transduction in Biomolecular Machines. Annu. Rev. Biophys. 2023, 52, 525–551.

(17) Li, C.; Voth, G. A. A quantitative paradigm for water-assisted proton transport through proteins and other confined spaces. Proc. Natl. Acad. Sci. USA 2021, 118, e2113141118.

(18) König, P.; Ghosh, N.; Hoffmann, M.; Elstner, M.; Tajkhorshid, E.; Frauenheim, T.; Cui, Q. Toward theoretical analyis of long-range proton transfer kinetics in biomolecular pumps. J. Phys. Chem. A 2006, 110, 548–563.

(19) Pomés, R.; Roux, B. Molecular mechanism of H+ conduction in the single-file water chain of the gramicidin channel. Biophys. J. 2002, 82, 2304–2316.

(20) Camilo, S. R. G.; Arantes, G. M. Flexibility and hydration of the Qo site determine multiple pathways for proton transfer in cytochrome bc1. bioRXiv:2024.08.22.609217 2024, under review in BBA Bioenergetics.

(21) Swainsbury, D. J. K.; Hawkings, F. R.; Martin, E. C.; Musia,l, S.; Salisbury, J. H.; Jackson, P. J.; Farmer, D. A.; Johnson, M. P.; Siebert, C. A.; Hitchcock, A.; Hunter, C. N. Cryo-EM structure of the four-subunit Rhodobacter sphaeroides cytochrome bc1 complex in styrene maleic acid nanodiscs. Proc. Natl. Acad. Sci. USA 2023, 120, e2217922120.

(22) Klusch, N.; Dreimann, M.; Senkler, J.; Rugen, N.; Kuhlbrandt, W.; Braun, H.-P. CryoEM structure of the respiratory I + III2 supercomplex from Arabidopsis thaliana at 2Å resolution. Nat. Plants 2023, 9, 142–156.

(23) Di Trani, J. M.; Gheorghita, A. A.; Turner, M.; Brzezinski, P.; Ádelroth, P.; Vahidi, S.; Howell, P. L.; Rubinstein, J. L. Structure of the bc1–cbb3 respiratory supercomplex from Pseudomonas aeruginosa. Proc. Natl. Acad. Sci. USA 2023, 120, e2307093120.

(24) Esser, L.; Elberry, M.; Zhou, F.; Yu, C.-A.; Yu, L.; Xia, D. Inhibitor-Complexed Structures of the Cytochrome bc1 from the Photosynthetic Bacterium Rhodobacter sphaeroides. J. Biol. Chem. 2008, 283, 2846–2857.

(25) Field, M. J. A Pratical Introduction to the Simulation of Molecular Systems, 1st ed.,, Cambridge University Press: Cambridge, 1999.

(26) Teixeira, M. H.; Curtolo, F.; Camilo, S. R. G.; Field, M. J.; Zheng, P.; Li, H.; Arantes, G. M. Modeling the Hydrolysis of Iron-Sulfur Clusters. J. Chem. Inf. Model. 2020, 60, 653–660.

(27) Arantes, G. M.; Ribeiro, M. C. C. A Microscopic View of Substitution Reactions Solvated by Ionic Liquids. J. Chem. Phys. 2008, 128, 114503.

(28) Stewart, J. J. P. Optimization of Parameters for Semiempirical Methods V: Modification of NDDO Approximations and Application to 70 Elements. J. Mol. Model. 2007, 13, 1173–1213.

(29) Galassi, V. V.; Arantes, G.M. Partition, Orientation and Mobility of Ubiquinones in a Lipid Bilayer. Biochim. Biophys. Acta 2015, 1847, 1345– 1573.

(30) Teixeira, M. H.; Arantes, G. M. Effects of Lipid Composition on Membrane Distribution and Permeability of Natural Quinones. RSC Adv. 2019, 9, 16892–16899.

(31) Huang, J.; Rauscher, S.; Nawrocki, G.; Ran, T.; Feig, M.; de Groot, B. L.; Grubmuller, H.; MacKerell Jr, A. D. CHARMM36m: an improved force field for folded and intrinsically disordered proteins. Nat. Methods 2017, 14, 71–73.

(32) Jorgensen, W. L.; Chandrasekhar, J.; Madura, J. D.; Impey, R. W.; Klein, M. L. Comparison of Simple Potential Functions for Simulating Liquid Water. J. Chem. Phys. 1983, 79, 926–935.

(33) Field, M. J. pDynamo3 Molecular Modeling and Simulation Program. J. Chem. Inf. Model. 2022, 62, 5849–5854.

(34) The pDynamo 3 molecular modeling and simulation program. https://github.com/pdynamo/pDynamo3, Accessed: 2024-11-20.

(35) Arantes, G. M. Dataset: Redox-Activated Proton Transfer through a Redundant Network in the Qo Site of Cytochrome bc1. 2024; 10.5281/zenodo.14198667, Accessed: 2024-11-20.

(36) Torrie, G. M.; Valleau, J. P. Nonphysical Sampling Distributions in Monte Carlo Free-Energy Estimation: Umbrella Sampling. J. Comp. Phys. 1977, 23, 187–199.

(37) Roux, B. The Calculation of the Potential of Mean Force Using Computer Simulations. Comp. Phys. Comm. 1995, 91, 275–282.

(38) Johnson, R. W. An Introduction to the Bootstrap. Teaching Statistics 2001, 23, 49–54.

(39) Miki, T.; Miki, M.; Orii, Y. Membrane potential-linked reversed electron transfer in the beef heart cytochrome bc1 complex reconstituted into potassium-loaded phospholipid vesicles. J. Biol. Chem. 1994, 269, 1827–1833.

(40) Osyczka, A.; Moser, C. C.; Daldal, F.; Dutton, P. L. Reversible redox energy coupling in electron transfer chains. Nature 2004, 427, 607– 612.

(41) Riccardi, D.; König, P.; Prat-Resina, X.; Yu, H.; Elstner, M.; Frauenheim, T.; Cui, Q. “Proton Holes” in Long-Range Proton Transfer Reactions in Solution and Enzymes: A Theoretical Analysis. J. Am. Chem. Soc. 2006, 128, 16302–16311.

(42) Borek, A.; Wójcik-Augustyn, A.; Kuleta, P.; Ekiert, R.; Osyczka, A. Identification of hydrogen bonding network for proton transfer at the quinol oxidation site of Rhodobacter capsulatus cytochrome bc1. J. Biol. Chem. 2023, 299, 105249.

(43) Hsueh, K.-L.; Westler, W. M.; Markley, J. L. NMR Investigations of the Rieske Protein from Thermus thermophilus Support a Coupled Proton and Electron Transfer Mechanism. J. Am. Chem. Soc. 2010, 132, 7908–7918.

(44) Zu, Y.; Couture, M. M.-J.; Kolling, D. R. J.; Crofts, A. R.; Eltis, L. D.; Fee, J. A.; Hirst, J. Reduction Potentials of Rieske Clusters: Importance of the Coupling between Oxidation State and Histidine Protonation State. Biochemistry 2003, 42, 12400–12408.

(45) Brasseur, G.; Saribas, A.; Daldal, F. A compilation of mutations located in the cytochrome b subunit of the bacterial and mitochondrial bc1 complex. Biochim. Biophys. Acta 1996, 1275, 61–69.

(46) Egawa, T.; Lee, H. J.; Ji, H.; Gennis, R. B.; Yeh, S.-R.; Rousseau, D. L. Identification of heme propionate vibrational modes in the resonance Raman spectra of cytochrome c oxidase. Anal. Biochem. 2009, 394, 141–143.

(47) Goyal, P.; Yang, S.; Cui, Q. Microscopic basis for kinetic gating in cytochrome c oxidase: insights from QM/MM analysis. Chem. Sci. 2014, 6, 826–841.

(48) Sezer, M.; Woelke, A.-L.; Knapp, E. W.; Schlesinger, R.; Mroginski, M. A.; Weidinger, I. M. Redox induced protonation of heme propionates in cytochrome c oxidase: Insights from surface enhanced resonance Raman spectroscopy and QM/MM calculations. Biochim. Biophys. Acta 2017, 1858, 103–108.

(49) Pagacz, J.; Borek, A.; Osyczka, A. ROS production by cytochrome bc1: Its mechanism as inferred from the effects of heme b cofactor mutants. Biochim. Biophys. Acta 2025, 1866, 149513.

(50) Gunner, M.; Amin, M.; Zhu, X.; Lu, J. Molecular mechanisms for generating transmembrane proton gradients. Biochim. Biophys. Acta 2013, 1827, 892–913.

(51) Saribas, A. S.; Ding, H.; Dutton, P. L.; Daldal, F. Tyrosine 147 of cytochrome b is required for efficient electron transfer at the ubihydroquinone oxidase site (Qo) of the cytochrome bc1 complex. Biochemistry 2005, 34, 16004–16012.

(52) Iverson, T. Catalytic mechanisms of complex II enzymes: A structural perspective. Biochim. Biophys. Acta 2013, 1827, 648–657.

(53) Pereira, C. S.; Teixeira, M. H.; Russell, D. A.; Hirst, J.; Arantes, G. M. Mechanism of rotenone binding to respiratory complex I depends on ligand flexibility. Sci. Rep. 2023, 13, 6738.

(54) Chung, I.; Wright, J. J.; Bridges, H. R.; Ivanov, B. S.; Biner, O.; Pereira, C. S.; Arantes, G. M.; Hirst, J. Cryo-EM structures define ubiquinone-10 binding to mitochondrial complex I and conformational transitions accompanying Q-site occupancy. Nat. Commun. 2022, 13, 2758.

(55) Das, D. K.; Medhi, O. K. The role of heme propionate in controlling the redox potential of heme: Square wave voltammetry of protoporphyrinato IX iron(III) in aqueous surfactant micelles. J. Inorg. Biochem. 1998, 70, 83–90.

