## Supporting Information file for "Redox-Activated Proton Transfer through a Redundant Network in the Q_*o*_ Site of Cytochrome *bc*_1_"

Guilherme M. Arantes*

*Department of Biochemistry, Instituto de Química, Universidade de São Paulo, Av. Prof. Lineu Prestes 748, 05508-900, São Paulo, SP, Brazil*

Contents:

- Figure S1: Comparison of potential energies obtained with different QC methods.
- Figure S2: Example of sampling and convergence of the computed free energy profiles.
- Table S1: Details of the three initial configurations used for QC/MM simulations.
- Supporting References.

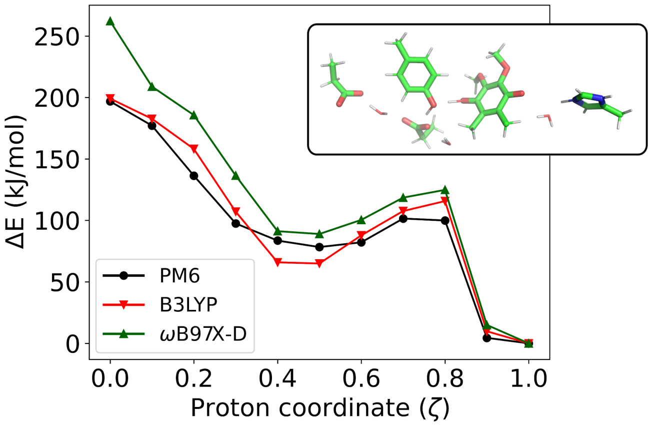


Figure S1: Comparison of relative potential energies (ΔE) obtained with three QC methods for the PT reaction of oxidized $QH^+$ along proton wire B (as in Fig. 4B). Geometries were optimized with the QC/MM potential in the $Q_o$ site environment (see Methods). The insert shows the optimized structure for $\zeta = 0.0$. Energies were calculated for the same geometries of the isolated QC region (in vacuum, a pure QC calculation) using PM6[28] and DFT functionals B3LYP[56] (with D3-corrections)[57] and $\omega$B97X-D,[58] both with the 6-31G(d,p) basis-set.[59] DFT calculations were done with the Gaussian09 program.[60] The average absolute difference between PM6 and B3LYP is just 7.6 kJ/mol and smaller than the difference between B3LYP and $\omega$B97X-D, two widely employed functionals with good accuracy.

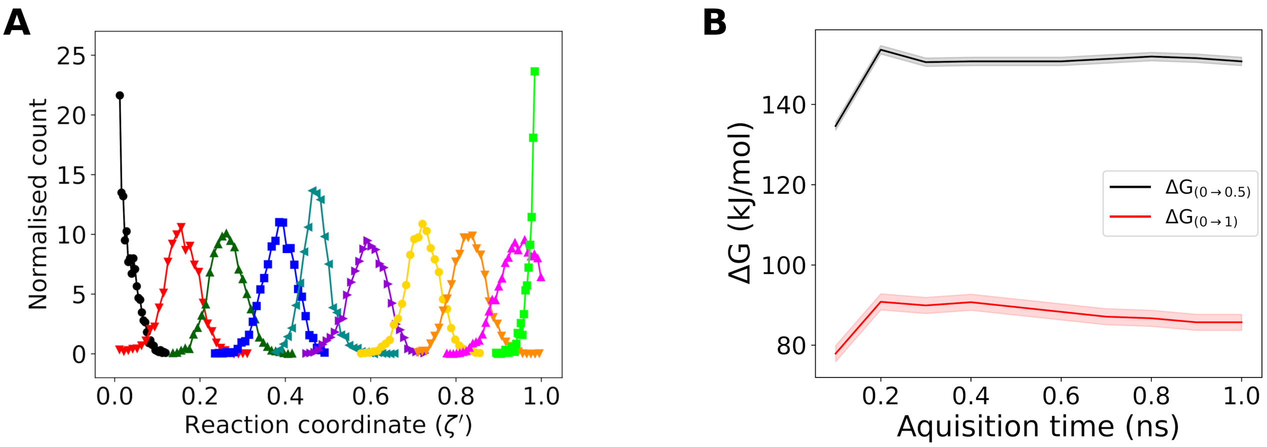


Figure S2: Example of sampling and convergence of the computed free energy profiles. **A**) Overlap of reaction coordinate ($\zeta'$ mCEC, eq. 1) during the US simulation with QC/MM potential (Fig. 6B) necessary for recovering the free energy profile. **B**) Convergence with simulation time of free energy differences (Fig. 6B) of the intermediate ($0 \rightarrow 0.5$, black) and the complete reaction ($0 \rightarrow 1$, red). The statistical error obtained from bootstrap analysis is shown with colored shadows. Variations in free energy differences are smaller than the statistical error after 0.5 ns, indicating convergence of the profiles.

Table S1: Conformers of side chains, atom-pair distances (in Å) and hydration (number of H-bonded waters) of reactive groups in the $Q_o$ site of the three initial configurations used for the QC/MM simulations.

| | Initial configuration | | |
|---|---|---|---|
| | 147A | 141D | 152D |
| Conformers: | | | |
| $Y147_{\chi1}$ | -g | t | t |
| $H276_{\chi1}$ | t | g | g |
| $D278_{\chi1}$ | -g | -g | -g |
| $E295_{\chi1}$ | -g | -g | -g |
| $E295_{\chi2}$ | -g | t | -g |
| $Y297_{\chi1}$ | g | t | t |
| Atom-pair distances: | | | |
| $O4_Q$-$N\delta_{H152}$ | 4.7 | 3.7 | 3.9 |
| $O1_Q$-$OH_{Y147}$ | 4.8 | 2.8 | 4.4 |
| $O1_Q$-$C\delta_{E295}$ | 7.2 | 4.7 | 5.8 |
| $O1_Q$-$C\gamma_{H276}$ | 12 | 8.1 | 7.5 |
| $OH_{Y147}$-$C\gamma_{H276}$ | 13 | 6.9 | 4.6 |
| $OH_{Y147}$-$C\delta_{E295}$ | 4.7 | 3.5 | 5.5 |
| $OH_{Y147}$-$OH_{Y297}$ | 2.8 | 13 | 5.5 |
| $C\delta_{E295}$-$C\gamma_{H276}$ | 8.6 | 4.7 | 4.8 |
| $C\delta_{E295}$-$C\gamma_{heme}$ | 6.4 | 6.5 | 10 |
| $OH_{Y297}$-$C\gamma_{heme}$ | 5.7 | 15 | 6.3 |
| $C\gamma_{H276}$-$C\gamma_{D278}$ | 4.7 | 5.8 | 5.7 |
| Hydration: | | | |
| $Q_{O4}$ | 2 | 1 | 1 |
| $Q_{O1}$ | 1 | 0 | 1 |
| $Y147_{OH}$ | 1 | 0 | 1 |
| $H276_N$ | 2 | 2 | 4 |
| $Heme_{O\gamma}$ | 5 | 3 | 3 |

Initial configurations were obtained from previous classical MD simulations and are representative of residue rotamers, atomic contacts and hydration of the two reactive binding modes identified for the Q-head.[20]

### Supporting References


(56) Becke, A. D. Density Functional Thermochemistry. III. The Role of Exact Exchange. *J. Chem. Phys.* **1993**, *98*, 5648.

(57) Grimme, S.; Hansen, A.; Brandenburg, J. G.; Bannwarth, C. Dispersion-Corrected Mean-Field Electronic Structure Methods. *Chem. Rev.* **2016**, *116*, 5105–5154.

(58) Chai, J.-D.; Head-Gordon, M. Systematic Optimization of Long-Range Corrected Hybrid Density Functionals. *J. Chem. Phys.* **2008**, *128*, 084106.

(59) Ditchfield, R.; Hehre, W.; Pople, J. A. Self-Consistent Molecular-Orbital Methods. IX. An Extended Gaussian-Type Basis for Molecular-Orbital Studies of Organic Molecules. *J. Chem. Phys.* **1971**, *54*, 724–728.

(60) Frisch, M. J. et al. Gaussian 09, Revision D.1. Gaussian, Inc., Wallingford CT, 2009.
